# Forced Abstinence from Alcohol Induces Sex-Specific Depression-like Behavioral and Neural Adaptations in Somatostatin Neurons in Cortical and Amygdalar Regions

**DOI:** 10.1101/2020.03.26.009514

**Authors:** Nigel C. Dao, Malini S. Nair, Sarah N. Magee, J. Brody Moyer, Veronica Sendao, Dakota F. Brockway, Nicole A. Crowley

## Abstract

Forced abstinence (FA) from alcohol has been shown to produce a variety of anxiety- and depression- like symptoms in animal models. Somatostatin (SST) neurons, a subtype of GABAergic neurons found throughout the brain, are a novel neural target with potential treatment implications in affective disorders, yet their role in alcohol use disorders (AUD) remains to be explored. Here, we examined the neuroadaptations of SST neurons during forced abstinence from voluntary alcohol consumption. Following six weeks of two-bottle choice alcohol consumption and protracted forced abstinence, male and female C57BL/6J mice exhibited a heightened, but sex-specific, depressive-like behavioral profile in the sucrose preference test (SPT) and forced swim test (FST), without changes in anxiety-like behaviors in the elevated plus maze (EPM) and open field test (OFT). FST-induced cFos expressions in the prefrontal cortex (PFC) and ventral bed nucleus of the stria terminalis (vBNST) were altered in FA-exposed female mice only, suggesting a sex-specific effect of forced abstinence on the neural response to acute stress. SST immunoreactivity in these regions was unaffected by forced abstinence, while differences were seen in SST/cFos co-expression in the vBNST. No differences in cFos or SST immunoreactivity were seen in the lateral central nucleus of the amygdala (CEA) and the basolateral amygdala (BLA). Additionally, SST neurons displayed opposing alterations in the PFC and vBNST, with heightened intrinsic excitability in the PFC and diminished intrinsic excitability in the vBNST. These findings provide an overall framework of forced abstinence-induced neuroadaptations in these key brain regions involved in emotional regulation and processing.

## INTRODUCTION

Alcohol use disorder (AUD) represents one of the most prevalent and costly neuropsychiatric disorders globally and domestically, costing the U.S. economy an estimated $249 billion due to losses in workplace productivity, and health care and criminal justice expenses (Sacks et al., 2015). Withdrawal from alcohol (both acute and protracted withdrawal) produces a host of negative emotional conditions, such as depression and anxiety (Becker, 2014; Hershon, 1977; Smith et al., 2019). These emotional states can increase the risk for relapse and further hamper an individual’s ability to abstain from alcohol. Importantly, the literature suggests that risk of relapse may be different across the sexes, with women more likely to relapse following negative affect-related situations, and men more likely to relapse when in the presence of other alcohol consumers (Peltier et al., 2019; White et al., 2015). Complementary research suggests that alleviating depressive symptoms following abstinence from alcohol may improve treatment outcomes for women (Annis et al., 1998; Holzhauer & Gamble, 2017). Taken together, these data point to a complex relationship between AUD, depression, and treatment outcome, likely moderated by sex.

Previous research has shown that forced abstinence (FA) from alcohol produces depressive-like behavior and a variety of neurobiological changes in rodent models. For instance, Vranjkovic et al. (2018) demonstrated that following six weeks of alcohol drinking, female C57BL/6J mice showed decreased time spent in the open arm of the elevated plus maze (EPM), and increased latency to feed in the novelty suppression of feeding test (NSFT), two commonly used models of anxiety-like behavior. In addition, this model produced increased immobility in the forced swim test (FST), though it should be noted males were not investigated in these studies (Holleran et al., 2016; Vranjkovic et al., 2018). Similarly, Valdez & Harshberger (2012) demonstrated that male Wistar rats exposed to chronic alcohol show increased immobility in the FST, and that this effect is further enhanced during protracted withdrawal.

AUD and major depressive disorder (MDD) are highly comorbid disorders with overlapping etiology. Novel fast acting antidepressants are able to reduce both binge drinking (Crowley et al., 2019) and depressive-like behavior following alcohol exposure (Holleran et al., 2016; Vranjkovic et al., 2018) further highlighting the potential overlapping neural circuitry involved in AUD and MDD. Importantly, both the clinical and preclinical depression literature has continuously pointed to a novel subpopulation of gamma aminobutyric acidergic (GABAergic) neurons as a protective, resiliency-conferring population. Somatostatin (SST) neurons are found throughout the brain, and have been implicated in a host of neuropsychiatric disorders in addition to MDD, such as bipolar disorder, anxiety disorder, and schizophrenia (Fee et al., 2017). Previous research has shown that global genetic upregulation of SST neuronal function reduces depression- and anxiety-like behaviors in male and female mice (Fuchs et al., 2017), and deficits in SST expression are seen in the amygdala of postmortem samples of MDD patients (Douillard-Guilloux et al., 2017). However, despite the clear correlation between SST neuronal markers and their function in MDD, both in the human and preclinical animal literature, thus far SST neurons have been poorly investigated in the context of AUD and depression-like phenotypes seen during withdrawal from alcohol.

The current study had two aims: first, to replicate previous behavioral models of forced abstinence from alcohol with both male and female mice. The second aim was to bridge the resiliency-like effect of SST neuronal function seen in the animal depression literature with the alcohol literature, in order to establish whether SST neurons within the prefrontal cortex (PFC), bed nucleus of the stria terminalis (BNST), basolateral amygdala (BLA) and lateral central amygdala (CEA), brain regions known for their role in MDD (Fee et al., 2017) and chronic alcohol (Pleil et al., 2015), are altered following forced abstinence from alcohol.

## MATERIALS AND METHODS

### Animals

Male and female C57BL/6J (stock #000664) were purchased from the Jackson Laboratory. Hemizygous female SST-IRES-Cre::Ai9 were generated from homozygous SST-IRES-Cre (stock #013044, Jackson Laboratory) and homozygous Ai9 (stock #007909, Jackson Laboratory) parents. All mice were at least 8-weeks old at the beginning of alcohol drinking. Mice were individually housed with *ad lib* access to food and water, and were maintained on a 12h reverse light/dark cycle (lights off at 7am) in temperature- and humidity-controlled vivarium for at least one week prior to alcohol exposure. Mouse weights were monitored weekly. All procedures were approved by the Pennsylvania State University’s Institutional Animal Care and Use Committee.

### Two-bottle choice (2BC) alcohol drinking

A schematic of the experimental timeline is displayed in **Figure 1A**. After one week of acclimatization to single housing, mice were randomly assigned to either drinking or non-drinking groups. Alcohol-drinking mice received continuous access to one sipper bottle of tap water and another of unsweetened, diluted alcohol. Alcohol concentrations increased from 3% in day 1 to 3, to 7% in day 3 to 9, to 10% in day 9 to 42. Bottles were weighed and refilled every 48-h. The positions of the bottles were randomized across mice, and within individual mice were switched weekly to avoid position bias. Following alcohol drinking, mice underwent forced abstinence where they had access to water only. Non-drinking control mice received continuous access to two water bottles only throughout the experiment. Escalation of alcohol drinking and forced abstinence were modeled off of Holleran & Winder (2016).

**Figure 1.**
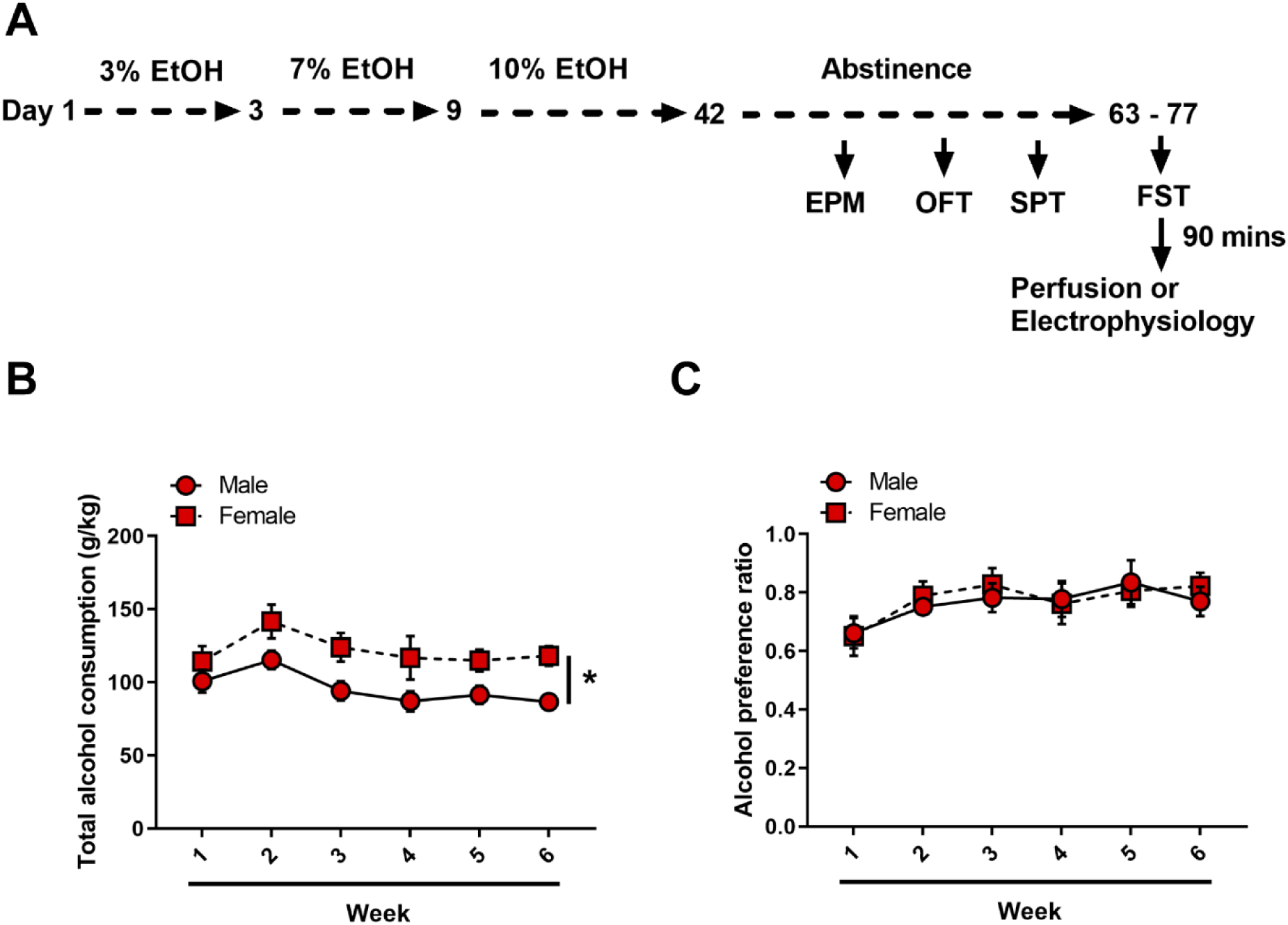
Forced abstinence from alcohol in alcohol ramp two-bottle choice paradigm. (A) Experimental timeline. Following 6 weeks of an alcohol ramp procedure, mice underwent a behavior battery (each test one week apart). Mice that were used in electrophysiology experiments did not undergo EPM, OPF or SPT. (B) Female mice consistently drink more than male mice. (C) Both sexes equally preferred alcohol over water. * *p* < 0.05

### Behavioral testing

Mice were tested for anxiety- and depression- like behaviors during abstinence, starting with elevated plus maze (EPM), open field test (OFT), sucrose preference test (SPT) and lastly forced swim test (FST). Prior to behavior, mice were brought to the testing room and allowed to rest for at least 30 minutes. All tests were done 3-h into the dark cycle, under red light (6 lux). Female SST-Ai9 mice that were used in electrophysiology experiments did not undergo EPM, OFT and SPT, and were tested in the FST under normal light.

### Elevated Plus Maze

One week after the onset of abstinence, mice underwent EPM where they were placed in the center square of the maze (35 × 5 × 40 cm), facing a closed arm (20 cm arm wall height, transparent Plexiglass and grey floor). Mice were allowed to explore the maze for 5 minutes. Sessions were recorded with EthoVision XT video tracking system (Noldus, Leesburg, VA). Total time spent in open arms and number of entries to open arms were automatically analyzed by EthoVision XT.

### Open Field Test (OFT)

Mice were placed in a corner of a black Plexiglass arena (50 × 50 × 20 cm) and allowed to explore for 20 minutes. Sessions were recorded with EthoVision XT, and total time spent in center zone and number of entries to center zones were automatically analyzed by EthoVison XT. Center zone was defined as a 12.5 × 12.5 cm area at the center of the arena.

### Sucrose Preference Test (SPT)

Mice received access to one bottle of tap water and another of 1% (w/v) sucrose solution for 12-h, starting 3-h into the dark cycle. Bottles were weighed before and after testing. Sucrose preference ratio was calculated as the volume of sucrose solution consumed divided by total volume of fluid consumed.

### Forced Swim Test (FST)

Mice were placed in a transparent 5-liter glass beaker filled with approximately 3-liters of water (20 ± 1°C) for 6 minutes. Total time spent immobile during the last 4 minutes was recorded and manually scored by a researcher blind to experimental conditions.

### Fluorescence Immunohistochemistry

Ninety minutes after FST, mice were deeply anesthetized with Avertin (250 mg/kg) and perfused transcardially with ice-cold phosphate buffered saline (PBS, pH 7.4) and 4% paraformaldehyde (PFA, pH 7.4). Brains were removed, post-fixed in PFA overnight and placed in 30% (w/v) sucrose solution until sunk. 50-μm free floating sections were sliced with a Compresstome vibrating microtome VF-300-0Z (Precisionary Instruments LLC, Greenville, NC) in a 1:3 series and stored in 30% sucrose/30% ethylene glycol cryoprotectant at -20°C until processed.

Prior to immunostaining, slices underwent antigen retrieval in 10 mM sodium citrate buffer (pH 6.0) at 80°C for 30 minutes. Slices were washed three times in PBS for 10 minutes each, and permeabilized in 0.5% Triton X-100 in PBS for 60 minutes. Nonspecific binding was blocked with 5% normal goat serum (NGS) in 0.1% Triton X-100 in PBS for 60 minutes. Slices were then incubated in a primary antibody cocktail, including rabbit anti-cFos (1:4000, Cell Signaling Technology, Danvers, MA) and rat anti-somatostatin (1:500, Millipore, Burlington, MA) in 2.5% NGS in 0.1% Triton X-100 in PBS for 48-h at 4°C. Slices were rinsed three times with PBS for 10 min each, and incubated in a fluorophore-tagged secondary antibody cocktail, including goat anti-rabbit Alexa Fluor 488 (1:500, Cell Signaling Technology, Danvers, MA) and goat anti-rat Cy3 (1:500, Millipore, Burlington, MA) for 4-h at room temperature. Slices were rinsed again three times with PBS, with the last step including DAPI (1:10,000), mounted on glass slides, air-dried and coverslipped with Immunomount (ThermoFisher, Waltham, MA). Images were obtained with an Olympus BX63 upright microscope (Center Valley, PA) under matched exposure settings. Four to eight images from both hemispheres were taken per region.

Total cFos counts, SST immunoreactivity (IR), and cFos+/SST+ counts were quantified by researchers blind to experimental conditions using ImageJ (National Institutes of Health, Bethesda, MD). For total cFos counts, region of interest (ROI) was delineated, and cFos+ nuclei were automatically quantified under matched criteria for size, circularity and intensity. Each ROI’s total Fos count was divided by the ROI’s area to give a cFos total density value (Smith et al., 2019). cFos+/SST+ nuclei count was manually quantified, and divided by the ROI’s area to give a cFos+/SST+ density value. SST IR was quantified as mean fluorescence intensity of the ROI (Pleil et al., 2015). At least 3 sections per region (PFC, dBNST, vBNST, lateral CeA, BLA) were quantified and averaged to obtain one value per mouse.

### Electrophysiology

Whole-cell current clamp recordings were conducted similarly to those previously published (Crowley et al., 2019; Crowley et al., 2016). Regions for electrophysiology were determined by cFos and SST results. Based on these results and behavioral results, only female mice were explored. The regions of interest (PFC and vBNST) were identified according to the Allen Mouse Brain Atlas. Following alcohol exposure and abstinence, mice underwent FST. Ninety minutes following FST, female SST-IRES-Cre::Ai9 mice were deeply anesthetized via inhaled isoflurane and rapidly decapitated. Brains were rapidly removed and processed according to the NMDG protective recovery method (Ting et al., 2018). Briefly, brains were immediately placed in ice-cold, oxygenated N-methyl-D-glucamine (NMDG)-HEPES aCSF containing the following, in mM: 92 NMDG, 2.5 KCl, 1.25 NaH2PO4, 30 NaHCO3, 20 HEPES, 25 glucose, 2 thiourea, 5 Na-ascorbate, 3 Na-pyruvate, 0.5 CaCl2·2H2O, and 10 MgSO4·7H2O (pH to 7.3–7.4). 300 μM coronal slices containing the PFC and the vBNST were prepared on a Compresstome vibrating microtome VF-300-0Z (Precisionary Instruments, Greenville, NC), and transferred to heated (31°C) NMDG-HEPES aCSF for a maximum of 10 min. Slices were then transferred to heated (31°C), oxygenated normal aCSF (in mM: 124 NaCl, 4.4 KCl, 2 CaCl2, 1.2 MgSO4, 1 NaH2PO4, 10.0 glucose, and 26.0 NaHCO3, pH 7.4, mOsm 300-310), where they were allowed to rest for at least 1-h before use. Finally, slices were moved to a submerged recording chamber (Warner Instruments, Hamden, CT) where they were continuously perfused with heated recording aCSF at a rate of 2 ml/min. Recording electrodes (3–6 MΩ) were pulled from thin-walled borosilicate glass capillaries with a Narishige P-100 Puller (Amityville, NY).

SST-expressing neurons were identified in SST-IRES-Cre::Ai9 mice via presence of tdTomato fluorescence under a 40x immersed objective with 565 nm LED excitation. Measurements of intrinsic excitability included resting membrane potential (RMP), rheobase (the minimum amount of current needed to elicit an action potential during a current ramp protocol), action potential threshold (the membrane potential at which the first action potential fired), and the number of action potentials fired during a voltage-current plot protocol (V-I plot) with increasing steps of depolarizing currents (0-200 pA, 10 pA per step). Hyperpolarizing currents (not shown) were included as a control. Experiments were performed at both RMP and at the standard holding potential of -70 mV. Electrodes were filled with a potassium gluconate-based (KGluc) intracellular recording solution (in mM: 135 K-Gluc, 5 NaCl, 2 MgCl2, 10 HEPES, 0.6 EGTA, 4 Na2ATP, and 0.4 Na2GTP, 287-290 mOsm, pH 7.35).

Signals were digitized at 10 kHz and filtered at 3 kHz using a Multiclamp 700B amplifier, and analyzed using Clampfit 10.7 software (Molecular Devices, Sunnyvale, CA). For all measures, recordings were performed in a maximum of two neurons per subregion, per mouse, and n values reported reflect the total number of neurons.

### Exclusion Criteria

One male mouse did not consume alcohol (average of 14.75 g/kg per week vs. group average of 108.12 g/kg per week). Since this was the first behavioral assay, this mouse was excluded from all analyses. For immunofluorescence quantification, problematic images (e.g., tears, out of focus, debris) were excluded prior to de-blinding. For electrophysiology recordings, cells that did not fire during the current ramp or V-I plot protocol were excluded. Statistical outliers, identified by Grubb’s test, were excluded from the data set.

### Statistical Analysis

Data was analyzed in Graphpad Prism 7.0 (San Diego, CA). All datasets were checked for normality (D’Agostino-Pearson’s test) and homogeneity (Barlett’s test). If found violated, nonparametric Kruskall-Wallis test was performed and Dunn’s multiple comparison was used as a posthoc test. Ordinary two-way ANOVA was used for all behavior and fluorescence immunohistochemistry data, followed by Tukey’s multiple comparison posthoc test. One sample *t* test was used to examine alcohol preference over water (theoretical mean 0.5). Student’s *t* test was used to analyze RMP, rheobase and action potential thresholds, while mixed-model two-way ANOVA was used to analyze SST V-I protocol. Statistical significance threshold was set at α = 0.05. Data presented show means and standard error of the mean (SEM).

## RESULTS

Male and female C57BL/6J mice underwent either an alcohol ramp two-bottle choice procedure and forced abstinence (N = 11 males and 10 females) or control condition (N = 12 males and 10 females) (**Figure 1A**). On average, over the six weeks of alcohol exposure, male mice consumed significantly less alcohol per body weight as compared to female mice (F_time_(5, 95) = 6.314, *p* < 0.001, F_sex_(1,19) = 7.060, *p* = 0.015, F_sex x time_(5,95) = 0.685, *p* = 0.635) (**Figure 1B**). Alcohol preference over water did not differ between male and female mice (F_time_(5, 95) = 5.839, *p* < 0.001, F_sex_(1,19) = 0.041, *p* = 0.84, F_sex x time_(5,95) = 0.535, *p* = 0.749), and both sexes displayed a strong preference for alcohol over water (male: one-sample *t*(10) = 5.391, *p* < 0.001, female: one-sample *t*(9) = 7.102, *p* < 0.001) (**Figure 1C**).

### Depressive-like behavioral phenotypes following forced abstinence from alcohol

Following the first week of abstinence, mice underwent multiple assays (one per week) to assess anxiety- and depression- like behavioral states. In the EPM, open arm duration was comparable between sexes and alcohol conditions (F_sex_(1, 39) = 0.379, *p* = 0.531, F_FA_(1, 39) = 0.991, *p* = 0.325, F_sex x FA_(1, 39) = 0.4, *p* = 0.530; **Figure 2A**). Females had higher frequency of entries into the open arms than males, (F_sex_(1, 39) = 5.275, *p* = 0.027) with no effects of FA conditions (F_FA_(1, 39) = 0.485, *p* = 0.489, F_sex x FA_(1, 39) = 0.044, *p* = 0.833; **Figure 2B**). In the OFT, no differences were seen in either open field center duration (F_sex_(1, 39) = 0.223, *p* = 0.639, F_FA_(1, 39) = 0.878, *p* = 0.354, F_sex x FA_(1, 39) = 0.066, *p* = 0.797; **Figure 2C**) or total distance travelled (F_sex_(1, 39) = 2.883, *p* = 0.097, F_FA_(1, 39) = 2.502, *p* = 0.121, F_sex x FA_(1, 39) = 1.575, *p* = 0.217; **Figure 2D**).

**Figure 2.**
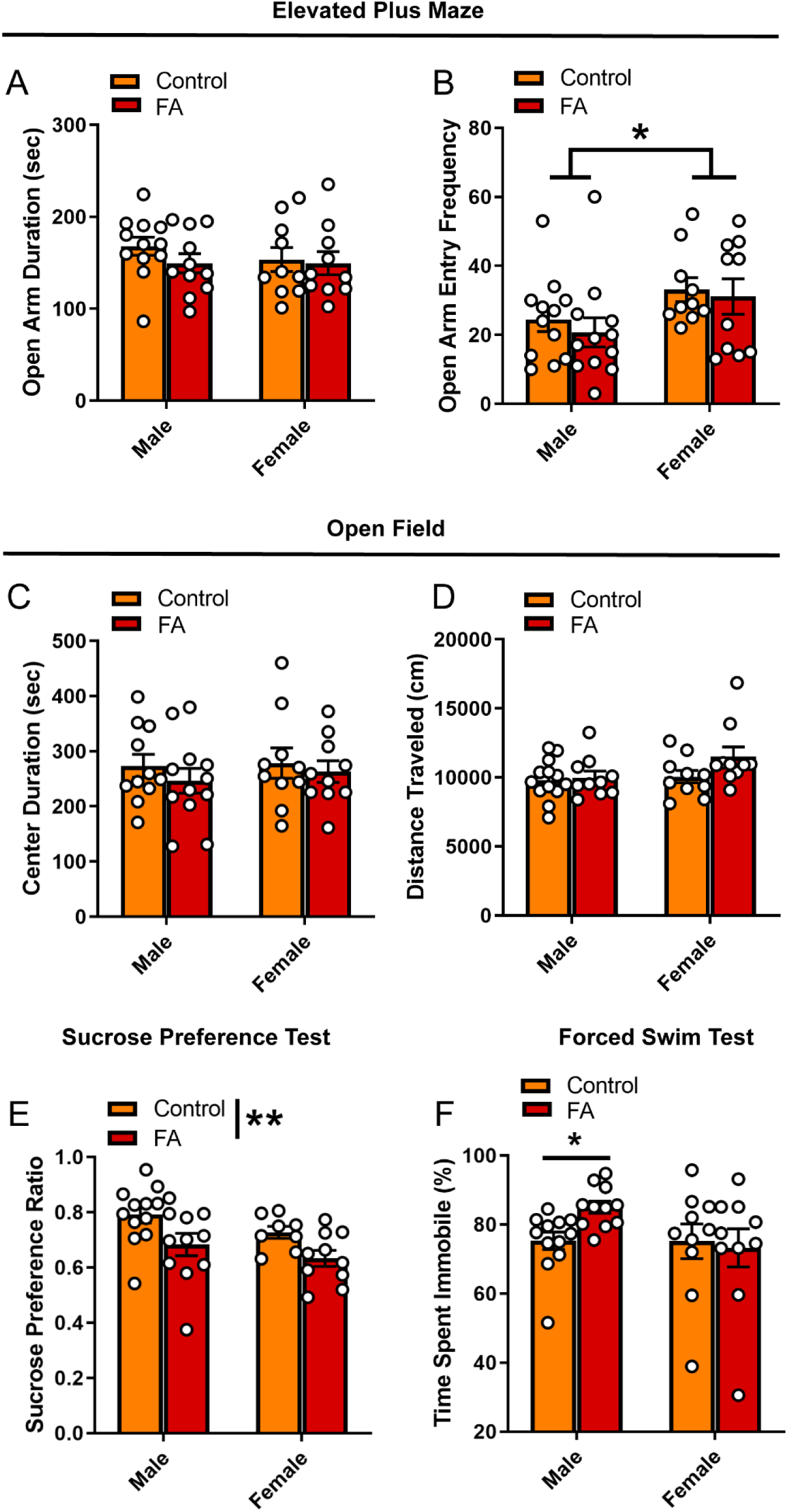
Forced abstinence from alcohol induces a depressive-like behavioral phenotype. (A) No significant difference in time spent in the open arm of the EPM was observed across sexes and FA conditions. (B) Female mice made more entries to the open arms than male mice, and this effect was not modulated by alcohol conditions. (C) No significant difference in time spent in the center zone of the OPT and (D) total distance travelled was observed across sexes and FA condition. (E) FA-exposed male and female mice displayed lower sucrose preference in the SPT than their control counterparts. (D) FA-exposed male mice spent more time immobile in the FST than control male mice. This effect was not observed in female mice. * *p* < 0.05. ** *p* < 0.01.

When probing depression-like behavior using the SPT and FST, key differences in both FA exposure and sex emerged. There was a significant main effect of FA history on sucrose preference (F_FA_(1, 37) = 9.388, *p* = 0.004), but no effect of sex (F_sex_(1, 37) = 2.983, *p* = 0.092) and no sex by FA interaction (F_sex x FA_(1, 37) = 0.049, *p* =0.825) (**Figure 2E**). Both male and female mice that underwent forced abstinence from FA showed lower sucrose preference, classically interpreted as an anhedonia phenotype. Interestingly, sex differences emerged in the FST. Though male mice exposed to FA showed a significant increase in immobility time, female mice did not (Kruskall-Wallis χ = 7.783, *p* = 0.05, male control vs. male EtOH: *p* = 0.02, female control vs. female EtOH: *p >* 0.99, Dunn’s posthoc test; **Figure 2F**). In sum, these data suggest that following forced abstinence from alcohol, male and female mice display a depressive-like behavioral profile without changes in anxiety-like behaviors.

### Forced swim stress-induced neuronal activation in the PFC is modulated by sex and forced abstinence from alcohol

Next, we probe the neural substrates that may underlie the changes in depressive-like behaviors in cortical and amygdalar areas, including the PFC, BNST, BLA and lateral CeA. In the PFC (**Figure 3A** for representative images), FST-induced neuronal activation, identified by expression of the immediately early gene marker cFos, showed significant main effect of sex (F_sex_(1, 33) = 22.44, *p* < 0.001) and significant interaction between sex and FA conditions (F_sex x FA_(1, 33) = 5.910, *p* = 0.020), without a main effect of FA (F_FA_(1, 33) = 1.198, *p* = 0.281; **Figure 3B**). Tukey’s posthoc test revealed that FA-exposed female mice showed significantly lower number of cFos nuclei in the PFC than control female mice (*p* < 0.05), control male mice (*p* < 0.001) and FA-exposed male mice (*p* < 0.001). Control female mice also had less cFos nuclei than FA-exposed male mice (*p* < 0.05). SST neuron-specific cFos expression (cFos+/SST+ nuclei) was not altered by sex and FA conditions (F_sex_(1, 36) = 2.166, *p* = 0.149, F_FA_(1, 36) = 2.140, *p* =0.152, F_sex x FA_(1, 36) = 0.202, *p* = 0.656; **Figure 3C**). There were also no changes in SST immunoreactivity (F_sex_(1, 35) = 1.889, *p* = 0.178, F_FA_(1, 35) = 0.082, *p* =0.775, F_sex x FA_(1, 35) = 0.026, *p* = 0.872; **Table 1**).

**Table 1.**
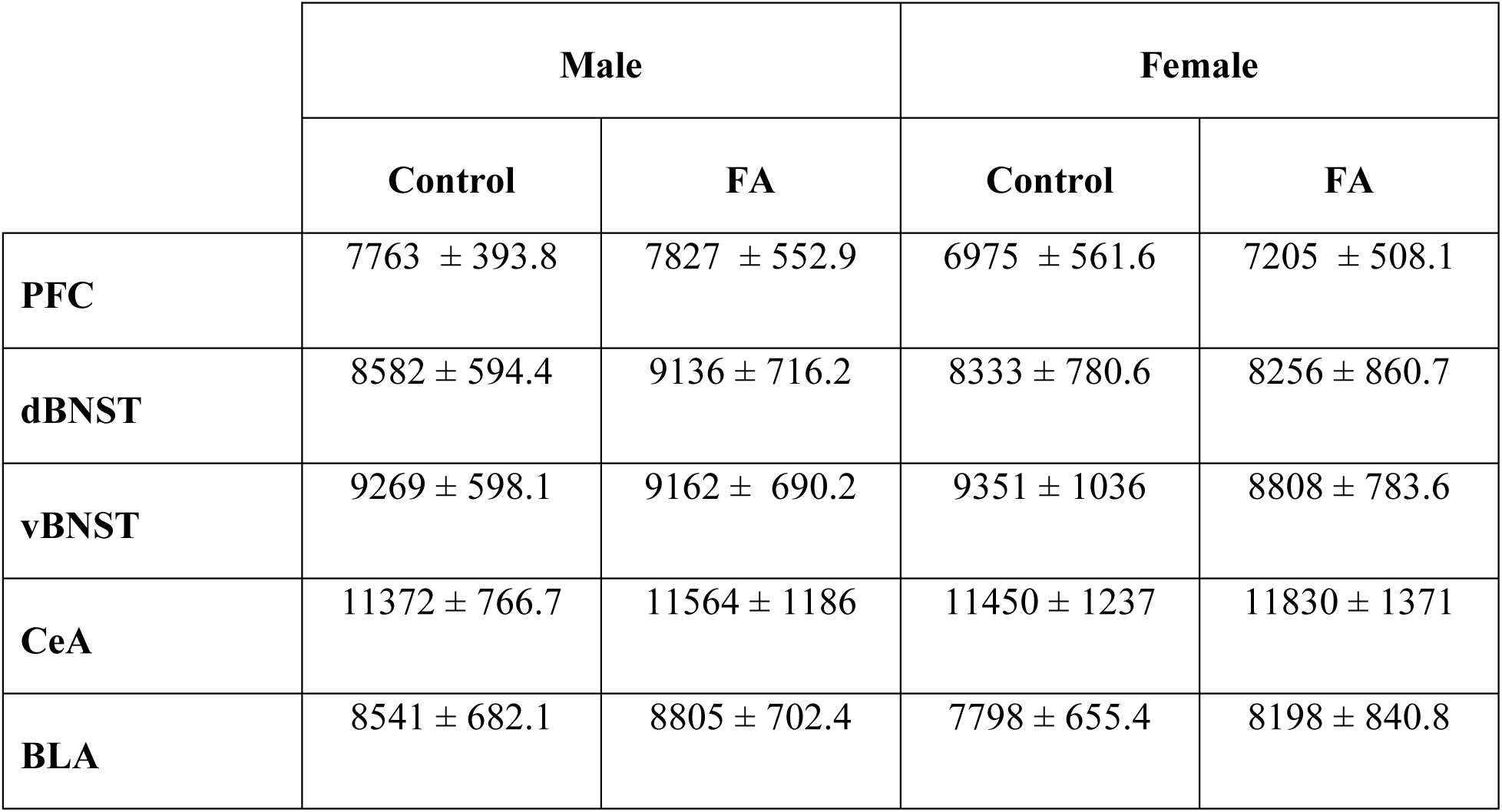
SST Immunoreactivity was not modulated by sex or forced abstinence from alcohol drinking.

**Figure 3.**
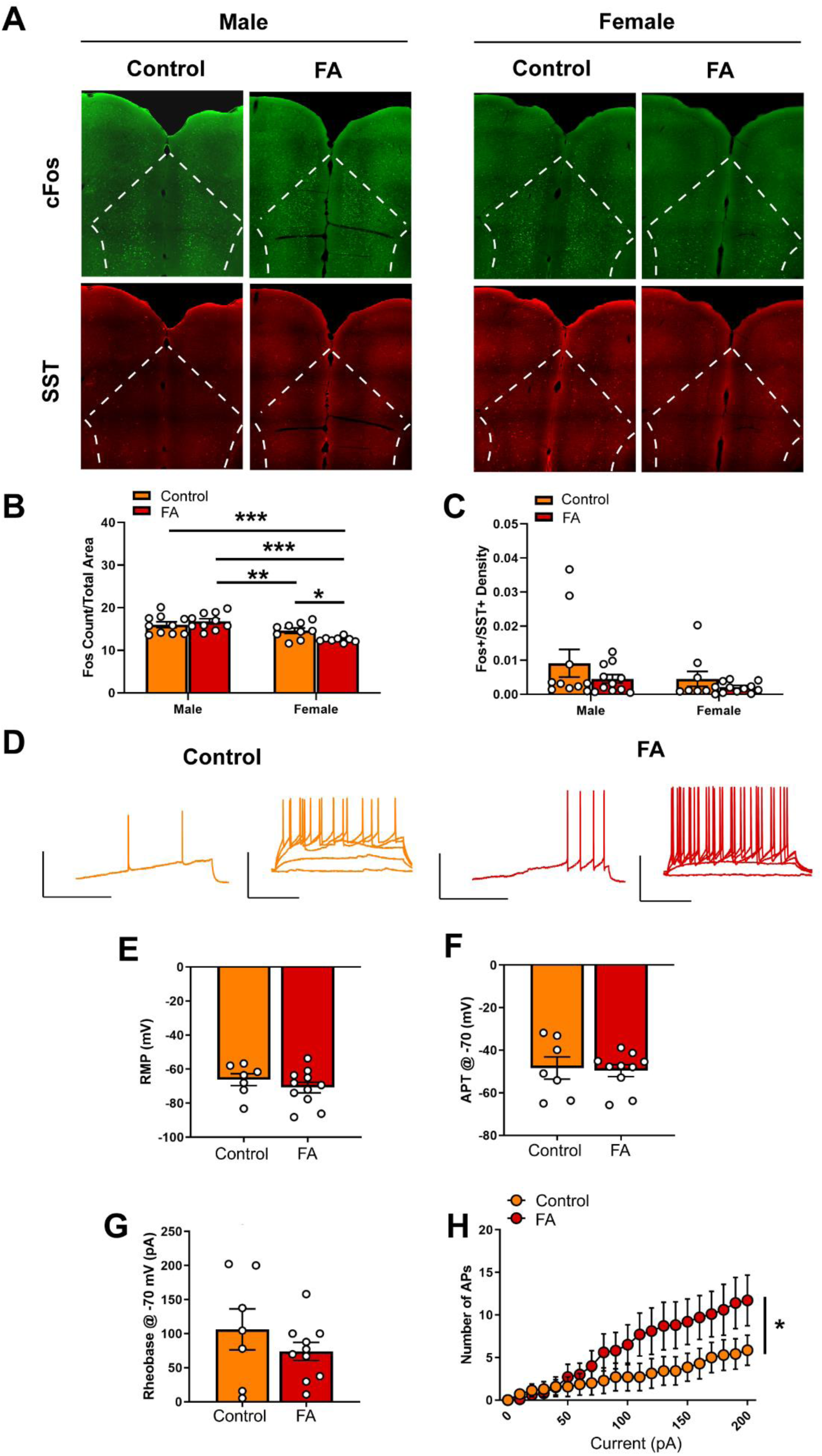
Forced abstinence from alcohol decreases forced swim stress-induced cFos expression and augments intrinsic excitability of SST neurons in the PFC of female mice. (A) Representative images of cFos (green) and SST (red) immunofluorescence with the PFC delineated for quantification. (B) Male mice displayed more cFos nuclei than female mice following forced swim, and FA-exposed female mice had less cFos nuclei than control female mice. (C) SST neuron-specific cFos expression was similar in both sexes and FA conditions. (D) Representative traces of current-injected firing from SST neurons in the PL during a ram*p* protocol (left) and V-I plot protocol (right) at -70 mV. Scale bars 50 mV × 200 ms. (E) RMP, (F) rheobase, (G) action potential threshold and (H) input resistance were unaltered in female PL SST neurons. (I) FA-exposed female SST neurons fired significantly more action potentials in the V-I plot protocol than control female SST neurons. * *p* < 0.05. ** *p* < 0.01. *** *p* < 0.001.

Given that only female mice displayed a change in FST-induced cFos expression in the PFC following forced abstinence from alcohol, we next performed whole-cell patch clamp electrophysiology in SST neurons in the PFC of female SST-Ai9 reporter (**Figure 3D** for representative recording traces). Current clamp experiments revealed that SST neurons had similar RMP (*t*(16) = 0.955, *p* = 0.353, **Figure 3E)**. When cells were held at -70 mV, action potential threshold (*t*(16) = 0.223, *p* = 0.825, **Figure 3F**) and rheobase (*t*(16) = 1.096, *p* = 0.290, **Figure 3G**) were unaltered between FA-exposed mice (n = 10 cells, N = 4 mice) and control mice (n = 7 cells, N = 3 mice). V-I plot at the holding potential of -70 mV indicated a significant main effect of current amplitude (F_current_(20, 300) = 15.257, *p* < 0.001) and a significant FA by current amplitude interaction (F_FA x current_(20, 300) = 2.970, *p* < 0.001), with no main effect of FA (F_FA_(1, 15) = 1.539, *p* = 0.234; **Figure 3H)**. SST neurons in the PFC of FA-exposed female mice fired significantly more action potentials in response to increasing steps of depolarizing currents than those in control female mice.

Recordings at RMP did not reveal any significant difference in rheobase (*t*(16) = 0.225, *p* = 0.824), action potential threshold (*t*(16) = 0.267, *p* = 0.792) and V-I plot (F_current_(20, 300) = 42.523, *p* < 0.001, F_FA_(1, 15) = 0.118, *p* = 0.735, F_FA x current_(20, 300) = 0.421, *p* = 0.987; data not shown) between FA-exposed mice and control mice.

Together, these data suggest that forced abstinence from alcohol dampened forced swim stress-induced neuronal activation in the female PFC, as indicated by reductions in cFos, likely via an increase in excitability of the GABAergic SST neurons.

### Forced swim stress-induced neuronal activation in the vBNST is modulated by sex and forced abstinence from alcohol

The dorsal BNST (**Figure 4A** for representative images) showed a significant main sex effect (F_sex_(1, 37) = 13.88, *p* < 0.001), but no main effect of FA (F_FA_(1, 37) = 1.646, *p* = 0.207) or sex by FA interaction (F_sex x FA_(1, 37) = 0.354, *p* = 0.149) in FST-induced cFos expression (**Figure 4B)**. Tukey’s posthoc test revealed that FA-exposed male mice had higher number of cFos nuclei than control female mice (*p* < 0.01) and FA-exposed female mice (*p* < 0.05). FA-exposed male and female mice had comparable number of cFos nuclei to their control counterparts (FA-exposed male vs. control male: *p* = 0.98, FA-exposed female vs. control female: *p* > 0.99). SST neuron-specific cFos expression (Fos+/SST+ nuclei) in the dorsal BNST was unaltered by sex and FA conditions (F_sex_(1, 38) = 0.178, *p* = 0.675, F_FA_(1, 38) = 0.134, *p* =0.715, F_sex x FA_(1, 38) = 0.0, *p* = 0.988; **Figure 4C**). SST immunoreactivity was intact across sexes and FA conditions (F_sex_(1, 37) = 0.106, *p* = 0.446, F_FA_(1, 37) = 0.106, *p* = 0.746, F_sex x FA_(1, 37) = 0.184, *p* = 0.669; **Table 1**).

**Figure 4.**
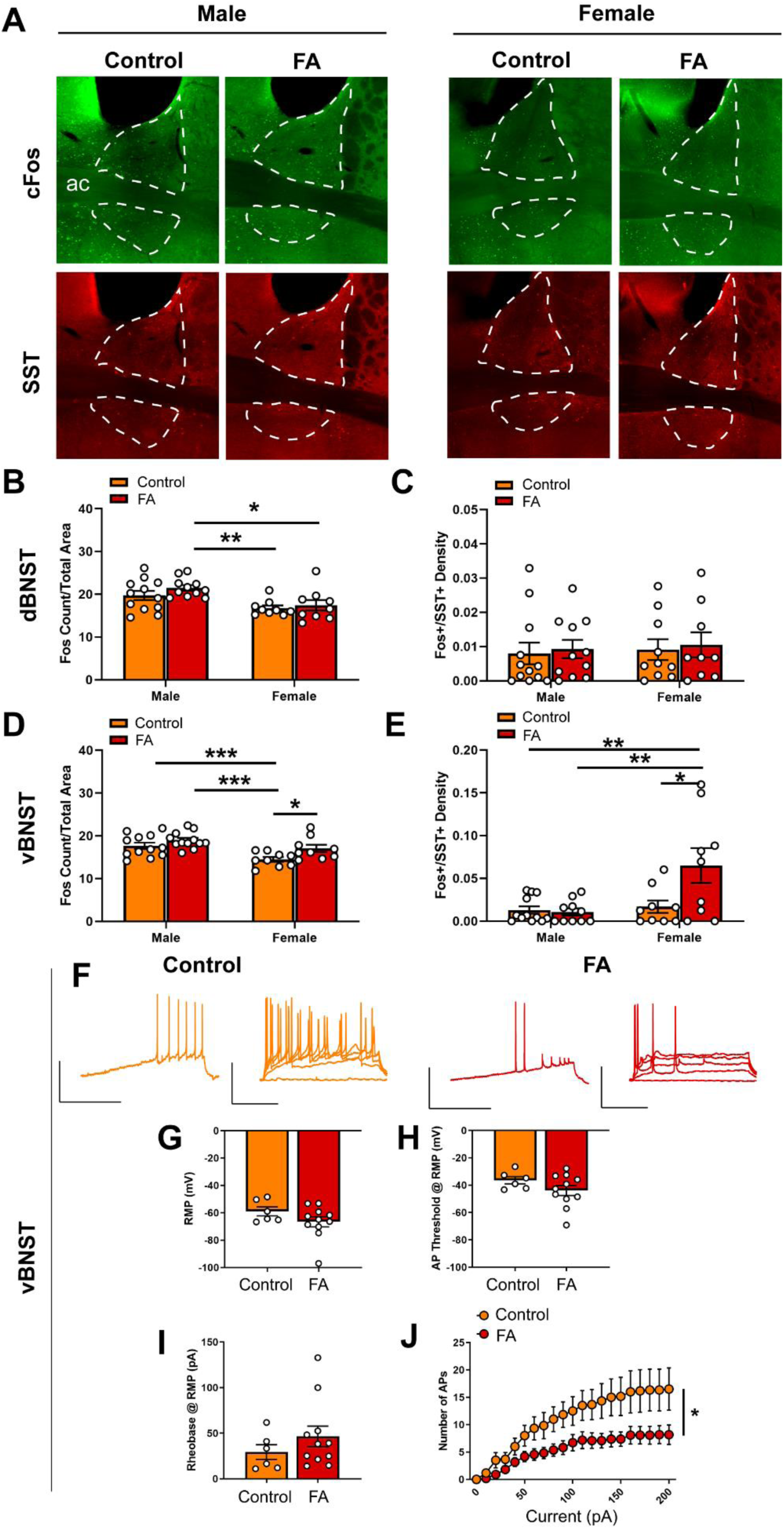
Forced abstinence from alcohol augments forced swim stress-induced neuronal activation and decreases intrinsic excitability of SST neurons in the vBNST. (A) Representative images of cFos (green) and SST (red) immunofluorescence in the BNST with the dorsal and ventral subregions delineated for quantification. (B) FA-exposed male mice had a higher number of cFos nuclei in the dorsal BNST than control female mice and FA-exposed female mice. (B) SST neuron-specific cFos expression in the dorsal BNST was comparable across sexes and FA conditions. (C) Control female mice had a lower number of cFos nuclei in the vBNST than FA-exposed female mice, control male mice and FA-exposed male mice. (D) FA-exposed female mice had a higher number of SST neuron-specific cFos nuclei than control female mice, control male mice and FA-exposed male mice. (F) Representative traces of current-injected firing from SST neurons in the vBNST during a ramp protocol (left) and V-I plot protocol (right) at RMP. (G) RMP, (H) rheobase and (I) action potential thresholds were unaltered in female vBNST SST neurons. (J) FA-exposed female SST neurons displayed lowered input resistance than control female SST neurons. (K) FA-exposed female SST neurons fired significantly less action potentials in the V-I plot protocol than control female SST neurons. * *p* < 0.05. ** *p* < 0.01. *** *p* < 0.001.

The vBNST (**Figure 4A** for representative images) showed a significant main sex effect (F_sex_(1, 38) = 13.59, *p* < 0.001), a significant main FA effect (F_FA_(1, 38) = 7.739, *p* = 0.008), without a sex by FA interaction (F_sex x FA_(1, 38) = 0.815, *p* = 0.372) in FST-induced cFos expression (**Figure 4D)**. Tukey’s posthoc test revealed that FA-exposed female mice had a higher number of cFos nuclei than control female mice (*p* < 0.05). Control male mice and FA-exposed male mice also have higher number of cFos nuclei than control female mice (*p*’s < 0.001). Additionally, cFos+/SST+ nuclei in the vBNST revealed a significant main sex effect (F_sex_(1, 36) = 7.905, *p* = 0.008), a significant main effect of FA (F_FA_(1, 36) = 4.944, *p* = 0.032) and a significant sex by FA interaction (F_sex x FA_(1, 36) = 5.918, *p* = 0.020; **Figure 4E**). Tukey’s posthoc test indicated that FA-exposed female mice showed higher number of Fos+/SST+ nuclei than control female mice (*p* < 0.05), control male mice (*p* < 0.01) and FA-exposed male mice (*p* < 0.01). SST immunoreactivity was similar across sexes and FA conditions (F_sex_(1, 37) = 0.030, *p* = 0.862, F_FA_(1, 37) = 0.174, *p* = 0.678, F_sex x FA_(1, 37) = 0.078, *p* = 0.780; **Table 1**).

Current clamp recordings from SST neurons in female vBNST (**Figure 4F** for representative recording traces) revealed that the RMP (*t*(15) = 1.371, *p* = 0.190; **Figure 4G**), action potential threshold at RM*P* (*t*(15) = 1.387, *p* = 0.185; **Figure 4H**), and rheobase at RMP (*t*(15) = 1.034, *p* = 0.317; **Figure 4I**) were unaltered in FA-exposed mice (n = 11 cells, N = 4 mice), compared to control mice (n = 6 cells, N = 3 mice). These neurons also fired significantly less action potentials in response to increasing steps of depolarizing currents at RMP, as revealed by a significant main effect of current amplitude (F_current_(20, 300) = 26.770, *p* < 0.001), a significant main effect of FA (F_FA_(1, 15) = 6.688, *p* = 0.021) and a significant current by FA interaction (F_current x FA_(20, 300) = 2.737, *p* < 0.001; **Figure 4J**). Similar effects were seen when cells were held at -70 mV, including rheobase (*t*(15) = 0.851, *p* = 0.408), action potential threshold (*t*(15) = 0.508, *p* = 0.618), and V-I plot (F_current_(20, 300) = 23.391, *p* < 0.001, F_FA_(1, 15) = 4.329, *p* = 0.05, F_current x FA_(20, 300) = 3.426, *p* < 0.001; data not shown).

In sum, these data suggest forced abstinence from alcohol induced robust neuroadaptations in the vBNST of female mice, with an increase in overall vBNST neuronal activation, as indicated by cFos, following forced swim stress and a decrease in excitability of the SST subpopulation of vBNST GABAergic neurons.

### Forced swim stress-induced neuronal activation in the BLA and lateral CeA is not modulated by sex or forced abstinence from alcohol

Contrary to the PFC and the vBNST, both the BLA and lateral CeA did not display any appreciable alteration in forced swim stress-induced cFos expression across sexes and FA conditions. In the BLA (**Figure 5A** for representative images), there was a significant main sex effect (F_sex_(1, 39) = 5.715, *p* = 0.021), but not main FA effect (F_FA_(1, 39) = 0.041, *p* = 0.839) or a sex by FA interaction (F_sex x FA_(1, 39) = 0.2850, *p* = 0.596; **Figure 5B**). In addition, Tukey’s posthoc test did not reveal any significant pair-wise difference. Consistent with the literature, very little SST immunoreactivity was observed in the BLA. Despite this, SST immunoreactivity was similar across sexes and FA conditions in the BLA (F_FA_(1, 39) = 0.870, *p* = 0.356, F_sex_(1, 39) = 0.211, *p* = 0.648, F_sex x FA_(1, 39) = 0.008, *p* = 0.925, **Table 1)**.

**Figure 5.**
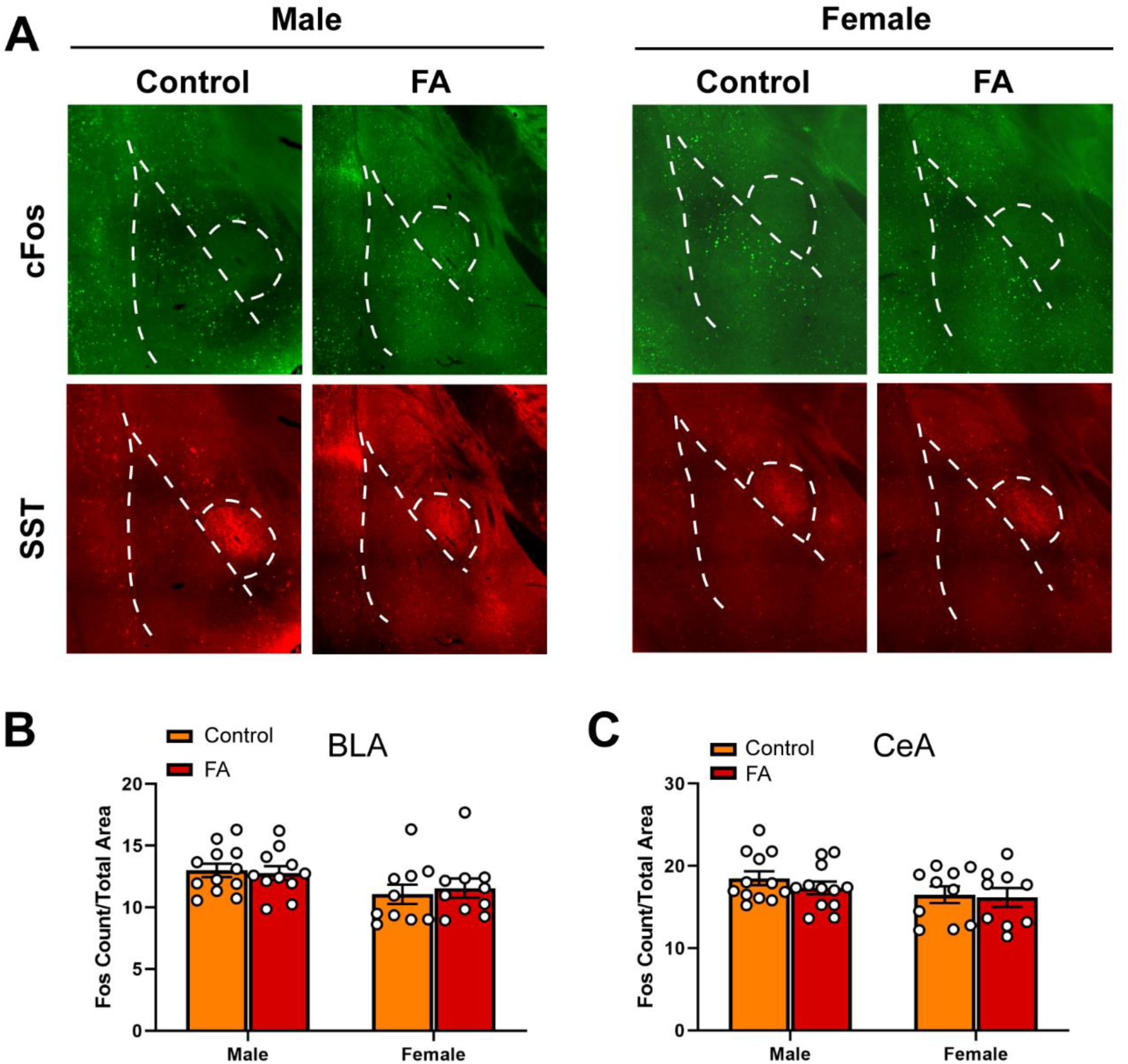
Forced abstinence from alcohol did not influence neuronal activation following forced swim stress in the BLA and CeA. (A) Representative images of cFos (green) and SST (red) immunofluorescence in the amygdala with the BLA and CeA delineated for quantification. (B) There was no significant difference in the number of cFos nuclei in the BLA between sexes and FA conditions. (C) There was no significant difference in the number of cFos nuclei in the CeA between sexes and FA conditions.

In the CeA (**Figure 5A**), there was no main sex effect (F_sex_(1, 39) = 2.836, *p* = 0.100), no main FA effect (F_FA_(1, 39) = 0.650, *p* = 0.424), and no sex by FA interaction (F_sex x FA_(1, 39) = 0.2147, *p* = 0.645; **Figure 5C**) in FST-induced cFos expression. SST immunoreactivity in the CeA was unaltered by FA or sex (F_FA_(1, 39) = 0.063, *p* = 0.802, F_sex_(1, 39) = 0.022, *p* = 0.881, F_sex x FA_(1, 39) = 0.006, *p* = 0.934, **Table 1)**. Because of this, electrophysiology was not conducted in the BLA and lateral CeA.

## DISCUSSION

The current study aimed to replicate previously published findings of abstinence-induced depression in both male and female rodents, and to understand the role SST neurons throughout the brain may play in this phenotype. Both male and female C57BL/6J mice showed an increased depressive-like behavioral profile following forced abstinence from alcohol, without aberrations in anxiety-like behaviors. Though both sexes showed a decrease in sucrose preference (a behavioral phenotype classically interpreted as anhedonia), only male mice showed an increased time spent immobile in the FST. Additionally, we observed plasticity in cortical and amygdalar areas of FA-exposed female mice in response to forced swim stress, including alterations in neuronal activation and intrinsic excitability of the GABAergic SST neurons in the PFC and vBNST. These findings highlight the efficacy of this alcohol exposure procedure to model protracted, forced abstinence-induced affective disturbances in both sexes, as well as shed insight into the circuit and cell type-specific adaptations.

Our results corroborate previous preclinical animal studies on sex differences in voluntary alcohol drinking (Almeida et al., 1998; Crowley et al., 2019; Peltier et al., 2019; Priddy et al., 2017), in which female mice consistently consumed more alcohol than male mice across 6 weeks of access, despite both sexes showing similar alcohol preference over water (**Figure 1B, C**). In humans, while women historically tend to consume less alcohol than men (Rehm et al., 2010), this gap is rapidly closing with substantial increases in the prevalence of alcohol use and binge drinking among women and not men (Grucza et al., 2018; White et al., 2015). Women are also more susceptible to development of alcohol-associated neuropsychiatric disorders, including MDD, anxiety, post-traumatic stress disorder (PTSD) and stress-mediated relapse in alcohol use (Grant et al., 2017; Peltier et al., 2019). A lack of observable differences in anxiety-like behaviors in the EPM and OFT between control and FA-exposed mice in our study could indicate a transition from the heightened anxious state associated with acute withdrawal and early abstinence (Lee et al., 2015; Pleil et al., 2015) to a more depressive-like state during protracted abstinence (Driessen, 2001; Heilig et al., 2010; Stevenson et al., 2009).

Previous studies using the same two-bottle choice paradigm demonstrated that female mice develop a heightened depressive-like behavioral phenotype during protracted abstinence from alcohol, as indicated by longer time spent immobile in the FST and longer latency to feed in the novelty-suppressed feeding test (NSFT), that can be alleviated by physical activity (Pang et al., 2013) or administration of the novel fast-acting antidepressant ketamine (Holleran et al., 2016; Vranjkovic et al., 2018). Interestingly, here we observed a more robust depressive-like behavioral phenotype in the FST in FA-exposed male mice, and not FA-exposed female mice. One hypothesis for these differences is that females transition to the abstinence-induced depressive-like state at a more rapid rate than males, such that heightened depressive-like behaviors in females may be observable at an earlier timepoint during abstinence. Alcohol-dependent men and women progress through courses of mood states at different rates during alcohol withdrawal (Bokström et al., 1991). Future experiments should assess the same depressive-like behavioral assays across multiple timepoints during protracted withdrawal. Another possibility is that male and female mice cope with acute stress and manifest depressive-like behaviors differently. Multiple studies in the literature (Colom-Lapetina et al., 2017, 2019; Kokras et al., 2015; Molendijk & de Kloet, 2019) have cautioned against over-interpretation of immobility duration in the FST, which classically has been interpreted as ‘behavioral despair’ or diminished motivation to escape a stressful environment (Porsolt et al., 1977). Recent work from Colom-Lapetina and colleagues (2017, 2019) suggests that female rats employ more active strategies to cope with the FST, including climbing and headshaking, while males employ the more classic strategy of immobility. Similar sex differences in coping strategy have also been found in conditioned fear response (Gruene et al., 2015). Furthermore, we observed a sex difference in the frequency of open arm entries in the EPM (**Figure 2B**), where more entries into the open arm is classically interpreted as lower level of anxiety. Overall, our results suggest that the behavioral adaptations in response to acute swim stress following forced abstinence may be more nuanced in female mice, and future studies should employ a more fine-tuned behavioral analysis to fully elucidate the affective perturbations associated with alcohol exposure and abstinence.

The corticolimbic circuit comprises of highly interconnected regions, including the medial prefrontal cortex (mPFC) and the amygdala and its subregions, that are critically involved in the regulation of emotional behaviors, stress response and reward seeking (Koob, 2009; Koob & Volkow, 2016). The mPFC, including the prelimbic (PL) and infralimbic (IL) subregions, receives glutamatergic inputs from the BLA that can promote anxiety-like behaviors (Felix-Ortiz et al., 2016) and are modulated by chronic stress (Lowery-Gionta et al., 2018; Marcus et al., 2019). Acute withdrawal from alcohol exposure has been found to disrupt synaptic transmission and intrinsic excitability of pyramidal neurons of the mPFC at different stages of alcohol dependence; particularly, the dependence-inducing chronic intermittent ethanol (CIE) model enhances excitatory drive and intrinsic excitability in pyramidal neurons (Cannady et al., 2018; Pleil et al., 2015), whereas the drinking-in-the-dark (DID) model of pre-dependence binge-like drinking impaired excitatory transmission (Crowley et al., 2019). Our FST-induced cFos expression data revealed that forced abstinence from alcohol dampened neuronal activation in the female PFC (**Figure 3B**). Hypoactivation of the mPFC in response to stress and subsequent dysregulation of amygdala activity have indeed been observed in humans with MDD and alcoholism (Covington et al., 2010; Johnstone et al., 2007; Seo et al., 2013), as well as rodents in chronic social defeat (Covington et al., 2010) and chronic unpredictable stress (Lam et al., 2018). Withdrawal from CIE exposure similarly reduces cFos expression in the PL and IL of male mice (Smith et al., 2019). Our model, which combines both stress (potential hypoactivation of PFC) and alcohol (potential hyperactivation) may result in more nuanced alterations in this region.

The downregulation of neuronal activation in the PFC may have resulted from augmented inhibitory tone from the GABAergic SST neurons, as we identified an increase in intrinsic excitability of these neurons (**Figure 3J, Figure 6)**. In the cortex, SST neurons are often postulated to be the gatekeeper of thalamo- and cortico-cortical excitatory inputs to pyramidal neurons, via their soma- and dendrite-targeting synapses (Urban-Ciecko & Barth, 2016). These neurons recently gained substantial interest for their role in a host of neuropsychiatric disorders, including MDD, bipolar disorder, schizophrenia (Abbas et al., 2018; Fee et al., 2017; Pantazopoulos et al., 2017). Global disinhibition of SST neurons via deletion of GABA_A_ receptors expressing the ϒ_2_ subunit produces an antidepressant-like behavioral and neural profile (Fuchs et al., 2017). Acute, chemogenetically induced inhibition of SST neurons in the mPFC is anxiogenic (Soumier & Sibille, 2014), pointing to an overall resiliency-conferring role of SST neurons in the cortex. In our study, it is unclear whether the hyperexcitability of SST neurons following forced abstinence from alcohol directly contributes to the depressive-like states, or is in fact a compensating mechanism to counteract the alcohol-induced adaptations. The latter interpretation, that SST hyperexcitability may be in response to alcohol-induced adaptations, is further supported by the opposing results seen in the vBNST. Future studies should further examine whether SST neurons in the PFC has a causal role in alcohol-related pathological behavioral adaptations.

**Figure 6.**
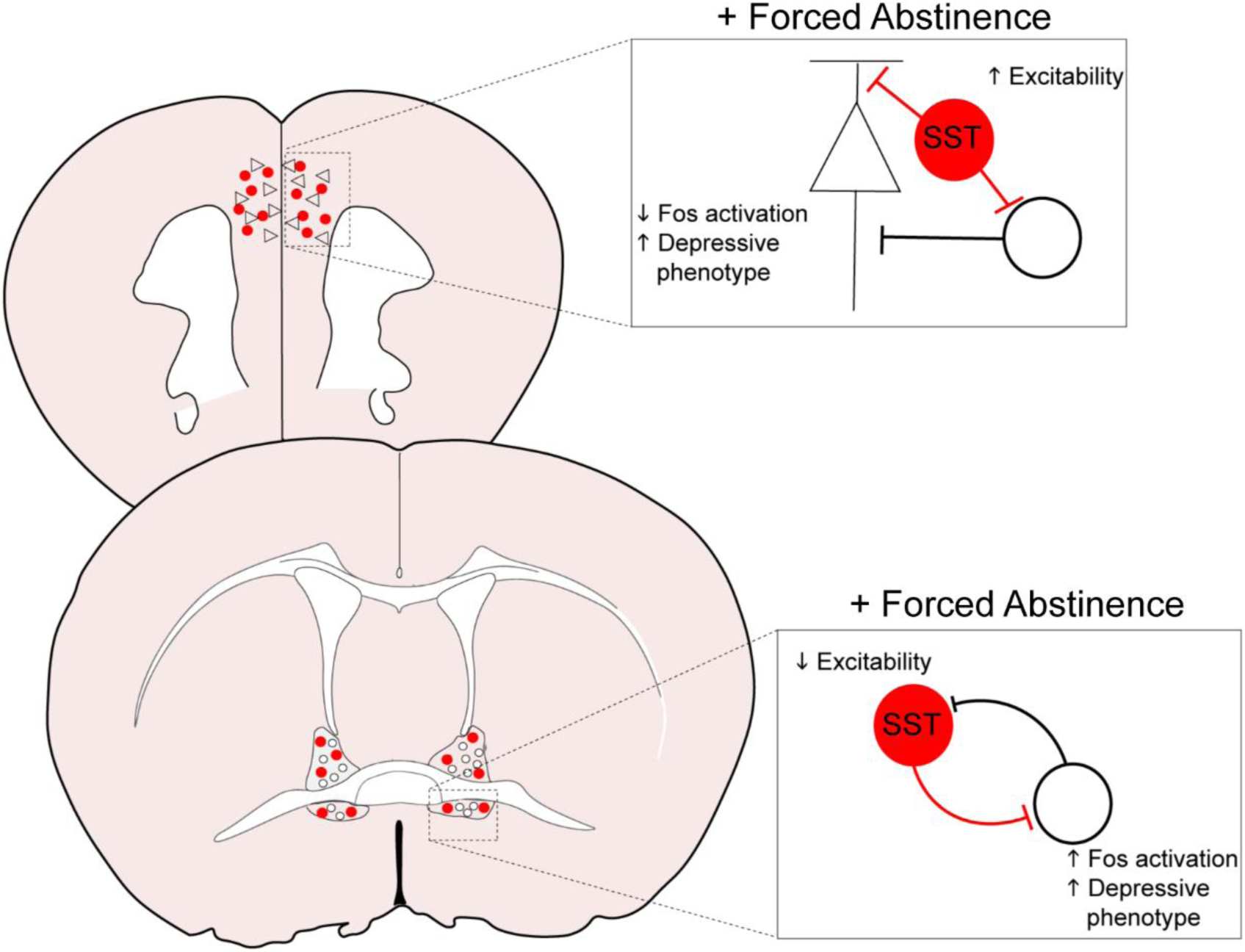
Overall model of neuroadaptations in cortical and amygdalar areas following forced abstinence from alcohol. In the PFC, forced abstinence-induced hyperexcitability of the GABAergic SST neurons may enhance inhibitory tone onto pyramidal neurons and other neighboring populations to dampen neuronal activation in response to forced swim stress. SST neurons in the vBNST otherwise showed diminished intrinsic excitability and consequently disinhibited the local network to enhance neuronal activation in response to forced swim stress.

The extended amygdala, including the BNST and the CeA, is critically involved in regulation of alcohol consumption and withdrawal-associated negative affect (Koob, 2009; Koob & Volkow, 2016; Torruella-Suárez et al., 2020; Vranjkovic et al., 2017). As in the mPFC, withdrawal from both the CIE and DID models modifies synaptic transmission and excitability in the extended amygdala, with heavy emphasis on hyperexcitability in the vBNST and enhanced inhibition of the CeA (Lee et al., 2015; Pleil et al., 2015; Roberto et al., 2010; Smith et al., 2019). In accordance with these results, our FST-induced cFos expression data demonstrated elevated neuronal activation in the vBNST of FA-exposed female mice (**Figure 4D**). SST neurons in the vBNST were also highly activated in response to FST in FA-exposed female, whereas this activation was very minimal in all other groups (**Figure 4E**). These data suggest that forced abstinence from alcohol may remodel the engagement of the multitude of the stress-responsive networks onto SST neurons in the vBNST (**Figure 6)**.

The circuit organization and functionality of these neuronal populations in the extended amygdala are not fully characterized. There is evidence suggesting that SST neurons in the lateral CeA directly synapse onto SST neurons in the oval nucleus of the BNST (Ahrens et al., 2018). Enhancing excitatory drive on the CeA SST neurons paradoxically disinhibits SST neurons in the oval nucleus in a kappa opioid receptor dependent manner to promote anxiety (Ahrens et al., 2018). As we did not observe any changes in anxiety-like behaviors, or any SST neuron-specific plasticity in the lateral CeA and the dorsal BNST, it is possible that the neuroadaptations in these anxiety-promoting populations may be transient and occur during acute withdrawal, subsiding before transition to a depressive-like state. The otherwise robust plasticity in the vBNST SST neurons may implicate a functional distinction between the dorsal and ventral subregions of the BNST. In addition to a dense population of corticotropin-releasing factor (CRF) neurons that are anxiogenic and alcohol drinking-promoting (Pleil et al., 2015), the vBNST includes an anxiolytic population of GABAergic neurons that project to the ventral tegmental area and the lateral hypothalamus (Marcinkiewcz et al., 2016; Pati et al., 2019). Withdrawal from CIE exposure augments excitability of CRF neurons and increases inhibition onto the anxiolytic midbrain-projecting neurons (Pati et al., 2019). Forced abstinence-induced decrease in intrinsic excitability of SST neurons in the vBNST may result in disinhibition of the CRF neurons to exacerbate negative affect and heighten the risk of stress-induced relapse. Nevertheless, there is no current evidence of direct synaptic input from SST neurons to CRF neurons or midbrain-projecting neurons in the vBNST. Future studies should examine the input and output pathways of these SST populations, as well as their functional role in regulation of alcohol drinking and emotional behaviors.

Notably, we observed a consistent sex effect in total cFos expression the PFC and the dorsal and ventral subregions of the BNST, in which male mice showed markedly higher level of FST-induced cFos expression (**Figure 3B, Figure 4B, 4D**). Given the fact that the majority of the previously cited studies on alcohol-induced neuroplasticity were performed in males, it is interesting to see that FA-exposed male mice seem to be relatively resilient against protracted forced abstinence-induced alterations. One possibility is that, since male mice consumed less alcohol than female mice in the two-bottle choice paradigm when normalized to body weight, the effect of forced abstinence from alcohol in male mice may be too subtle to be detected by the methods employed here. Additionally, as discussed with the sex differences in coping strategies in response to forced swim stress, passive coping in male mice may engage in a different circuit and cell-type specificity that could be overlooked. These observations further highlight the importance of sex as a biological variate in neurobiological studies in order to have a better understanding of the pathophysiology underlying neuropsychiatric disorders.

We did not observe any changes in SST protein expression in any cortical or amygdalar regions. We have previously reported that SST neurons in the PL cortex release SST peptide tonically and phasically in a frequency-dependent manner (Dao et al., 2019), hence altered excitability of these neurons in the PFC and the BNST may affect their capacity for neuropeptide release. The SST peptide acts on a family of G protein-coupled receptors, mainly inhibitory through Gi/o-dependent signaling (Barnett, 2003). However, SST receptor signaling in these regions has not been characterized. Future studies may merit from examination of the modulatory action of SST peptide, in healthy states and alcohol abuse-associated conditions.

## CONCLUSION

The current study identified a host of sex-dependent behavioral adaptations and neuroplasticity in corticolimbic regions that may underlie forced abstinence-induced affective perturbations. SST neurons in the PFC and vBNST emerge as a strong neural candidate that undergo robust plasticity following alcohol exposure and protracted abstinence. These results shed new insight into the neurobiological bases of highly comorbid neuropsychiatric diseases, including MDD and AUDs, and may aid in the development of new and effective treatments.

## ACKNOWLEDGEMENTS

The experiments were funded by The Brain and Behavior Research Foundation (NARSAD Young Investigator Award; NAC) and Penn State’s Social Science Research Institute (Level 2 Award). NCD and NAC conceived of and designed the experiments, NCD, MSN, SNM, JBM, VS and DFB conducted the experiments, NCD and NAC analyzed and interpreted the experiments, and NCD and NAC wrote the paper with feedback from all authors. The authors would like to thank Dr. Janine Kwapis (Department of Biology, Pennsylvania State University) for assistance in imaging.

## DISCLOSURES

The authors have no financial disclosures and no conflicts of interest to report.

